# Ultra-Accurate Classification and Discovery of Functional Protein-Coding Genes from Microbiomes Using FunGeneTyper: An Expandable Deep Learning-Based Framework

**DOI:** 10.1101/2022.12.28.522150

**Authors:** Guoqing Zhang, Hui Wang, Zhiguo Zhang, Lu Zhang, Guibing Guo, Jian Yang, Fajie Yuan, Feng Ju

## Abstract

High-throughput DNA sequencing technologies open the gate to tremendous (meta)genomic data from yet-to-be-explored microbial dark matter. However, accurately assigning protein functions to new gene sequences remains challenging. To this end, we developed FunGeneTyper, an expandable deep learning-based framework with models, structured databases and tools for ultra-accurate (>0.99) and fine-grained classification and discovery of antibiotic resistance genes (ARGs) and virulence factor or toxin genes. Specifically, this new framework achieves superior performance in discovering new ARGs from human gut (accuracy: 0.8512; and F1-score: 0.6948), wastewater (0.7273; 0.6072), and soil (0.8269; 0.5445) samples, beating the state-of-the-art bioinformatics tools and protein sequence-based (F1-score: 0.0556-0.5065) and domain-based (F1-score: 0.2630-0.5224) alignment approaches. We empowered the generalized application of the framework by implementing a lightweight, privacy-preserving and plug-and-play neural network module shareable among global developers and users. The FunGeneTyper^*^ is released to promote the monitoring of key functional genes and discovery of precious enzymatic resources from diverse microbiomes.

## Main

High-throughput DNA sequencing and metagenomics have generated extensive protein-coding gene (PCG) sequences from diverse environmental and human microbiomes^1–3^. Accurate classification of genes into related protein functions is the key to effective gene discovery. However, these datasets pose significant computational challenges in metagenomic studies. Sequence alignment (SA), implemented using NCBI’s BLAST^4^, usearch^5^, and Diamond^6^, is commonly used for functional annotation of PCGs^7^. To minimize false-positives, SA-based methods are routinely conducted with strict user-defined cutoffs or thresholds (alignment identity, coverage, and bit scores) to retain high-confidence best hits for each query sequence from validated databases. This practice is widely implemented in the development of tools for categorizing genes, including antibiotic resistance genes (ARGs)^8, 9^ and virulence factor genes (VFGs)^10^. SA-based approaches effectively predict functions between genes that share high homology (>80% identity ^8, 9^), but exclude distantly homogeneous genes that fall below arbitrarily-defined and one-size-fits-all cutoffs that may represent the majority of targeted functional genes in environmental samples (e.g. core ARGs in activated sludge^11^ and soil^12^). Therefore, these SA approaches with stringent bioinformatic cutoffs unavoidably generate numerous false-negative results and heavily underestimate true novelties and diversity of functional genes in largely uncultured bacteria, thus biasing research outcomes or conclusions. Therefore, it is crucial to develop intelligent and accurate classification paradigm and bioinformatic tools to overcome limitations of existing SA-based classification approaches. Importantly, this endeavor will accelerate discovery of new genes in future metagenomic-based environmental and human microbiome studies^13, 14^.

Hidden Markov models (HMM) with manual-crafted sequence alignments and scoring functions are powerful tools for protein domain-based functional gene annotation for detecting remote gene homologues with low sequence identity (< 30%) to known proteins^15, 16^. However, these methods rely on token (amino acid) matching, which fail to detect high-level semantic representation similarity or structure-level representation similarity, leading to false-positives^17^, and thus cannot distinguish functions of proteins in the same family^18^. In contrast, deep learning (DL) methods excel at learning rich and high-level semantic representations when sufficient training data are available, and are effective at identifying proteins with structural and functional similarities^19–22^. Specifically, ground-breaking big language models initially developed for natural language processing tasks have been successfully applied to protein function prediction tasks^23, 24^, often termed protein language models (PLMs). The high-level semantic representations learned from PLMs establish valid connections between sequences and function^25, 26^. Notwithstanding the power of PLMs, gene classification tasks, particularly identifying fine-grained protein function subclasses, pose challenges for data-hungry deep learning paradigms because of limited supervised training dataset for certain genes. Additionally, it remains unclear whether advanced PLMs perform better than state-of-the-art metagenomic bioinformatics tools at microbiome gene classification and discovery.

Here, we propose FunGeneTyper, a PLM-based deep-learning framework for accurate and expandable prediction of PCG function. FunGeneTyper implements a two-stage pipeline that separately handles the assignment of the main types and subtypes of PCG functional classes, reducing issues associated with insufficient training data during subtype-level predictions. To improve conciseness, it first performs standard classification of genes of the main types and then performs fine-grained retrieval by comparing similarities between learned protein subtype representations. FunGeneTyper models classify ARGs with ultra-high accuracy (>0.99) and outperforms the state-of-the-art SA and HMM-based methods and tools. Furthermore, we also demonstrate the generalized application of FunGeneTyper models in ultra-high classification of VFGs and introduce the adapter module, a lightweight neural network that can be inserted into the current backbone architecture to realize parameter-efficient training. The adapter-tuning-based FunGeneTyper models are expandable to the classification of various categories of genes and enables sharing of both task-agnostic and task-specific parameters without accessing the private training dataset. Thus, FunGeneTyper offers a unified and innovative way of integrating the global efforts of the microbiome and bioinformatics communities, endowing the FunGeneTyper framework with the ability to conduct unlimited prediction of functional gene categories beyond the ARGs and VFGs demonstrated here, which is key to accelerating the global discovery of new and precious genetic and enzymatic resources from microbiomes.

## Results

### FunGeneTyper framework, structured database, and deep learning models

FunGeneTyper is the first unified framework that utilizes DL models and structured functional gene datasets (SFGD) to develop new DL-based classifiers for any gene category via transfer learning. This novel framework achieves highly accurate PCG classification from metagenomic studies and extends the models to efficiently predict broad categories of gene functions from large varieties of microbiomes with corresponding customizable SFGD.

### Structured functional gene datasets

We deployed a transferable strategy to collect high-quality reference gene sequences to meet FunGeneTyper’s training requirements with high reliability (Fig. 1a). Experimentally-confirmed reference sequences of target genes from literature and/or expert-curated databases were used as the core dataset, and highly homologous protein sequences (at least 80% identity and 80% coverage) were extracted from Uniref100 database and used as the expanded functional genes dataset. A non-target sequence dataset was selected from Swiss-Prot database (version: June 2021) by excluding perfect matches to the target genes, and used as the negative training set so that FunGeneTyper could learn sufficient features of non-target genes. Core and expanded functional gene datasets and the non-target dataset were integrated to form the SFGD, which was organized hierarchically into a secondary structure based on gene ontology. The SFGD was divided into training, validation, and testing sets (ratio 6:2:2) and used to train the following two DL models.

**Fig. 1.**
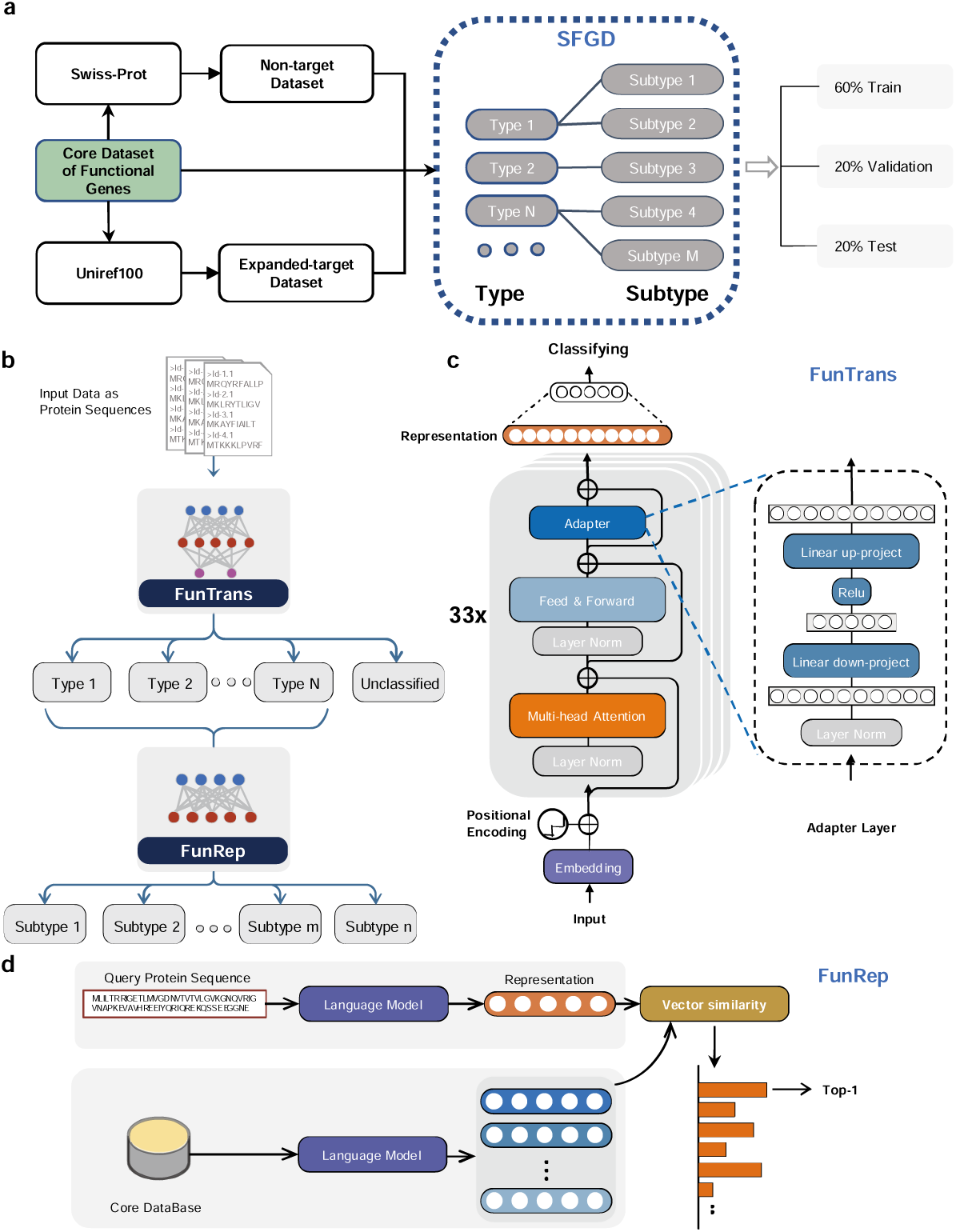
FunGeneTyper model design and database construction workflows. **a**, Process of preparing a structured functional gene dataset (SFGD). The data set is divided into the training set, validation set and testing set in a 6:2:2 ratio. **b**, Two-level hierarchical structure of FunGeneTyper. **c**, Schematic representation of FunTrans model. **d**, Schematic representation of FunRep model.

### Deep learning models

The framework has a top-down protein function prediction workflow featuring two DL models (Fig. 1b), FunTrans and FunRep, which progressively classify protein sequences from the upper (type) to lower (subtype) functional levels. FunGeneTyper was pre-trained on ESM-1b, a large-scale pre-trained protein sequence model based on the transformer architecture released by Facebook^22^. ESM-1b is composed of the 33-layer transformer architecture consisting of 650 million parameters trained on Uniref50. It has the superior capacity to infer fundamental structural and functional characteristics of proteins from sequences that can significantly increase the performance metrics for sequence-function tasks. Despite sharing the 33-layer transformer architecture, FunTrans and FunRep were independently constructed, trained, and optimized, to complete two-level functional classification tasks that successively assigned a PCG to its best matches of functional type and subtype in a structured database, respectively.

FunTrans distinguishes protein sequences and classify them into specific functional types equivalent to gene families with the same or similar functions. It is inspired by the fact that proteins with similar structures and functions clustered closer together in the embedding space. The main FunTrans structure is a 33-layer transformer that implements initial classification of input data (Fig. 1c). Adapter modules are inserted into each transformer block as trainable parameters. The adapter enables efficient fine-tuning of parameters for different gene classification tasks. High parameter sharing is achieved under the premise that the parameters of the original network remain unchanged. Adapter modules enable flexible and parameter-efficient transfer learning and prevent overfitting^27, 28^. FunTrans adds a nonlinear classification layer at the end of the sequence semantic representation for functional classification.

FunRep has a structure similar to that of FunTrans and can be used to embed representation retrieval for further subtype classification of protein function (Fig. 1d). It uses embedding representation retrieval to accurately predict functional subtypes of the FunTrans output results for classification. FunRep also adds an adapter layer to increase robustness and insight into a broader range of gene classification.

### FunGeneTyper classification performance and learning ability

The spread of antibiotic resistance has raised public health concerns globally^29^. Reliable ARG model classification is important for surveillance and control of the spread of antibiotic resistance, and achieving sufficient model sensitivity to remote homologues is key to discovering new ARGs. Therefore, we first classified ARGs and demonstrated the ability of the FunGeneTyper framework to achieve this goal. Before building the ARGs classification models, we constructed a hierarchical structured ARG database (SARD) based on antibiotic resistance ontology of the comprehensive antibiotic resistance database (CARD)^7^. Based on CARD’s ontological rules, ARGs were assigned to class and group hierarchies based on the types of drugs to which they confer resistance, and the subtypes of genes with the same resistance function, respectively (Dataset S1). To test and improve the sensitivity of the model, we used different identity thresholds to collect four non-target sequence sets from Swiss-Prot database —excluding ARGs — as negative training datasets, for model training (Supplementary Figure 1, see Methods). The addition of a negative training set allows the model to learn features of non-targeted genes, which gives the model the ability to directly classify targeted (e.g., ARGs) and non-targeted genes (e.g., non-ARGs) from new datasets to be tested. We evaluated the impact of four identity thresholds of the negative datasets on the learning features of the model. The results of five-fold cross-validation revealed that the model with 0% identity as the threshold for recruiting non-target sequences had the best performance metrics, including accuracy, recall, precision, and F1-score (Fig. 2a). Under these optimized conditions, the positive SARD set contained 61874 ARG sequences, including 2972 experimentally-confirmed core sequences inherited from the CARD and 58902 homology-predicted (>80% identity and >80% coverage) expanded ARG sequences from Uniref100. All ARG reference sequences were hierarchically assigned to 19 classes and 2972 groups (Dataset S2 and Supplementary Figure 2).

**Fig. 2.**
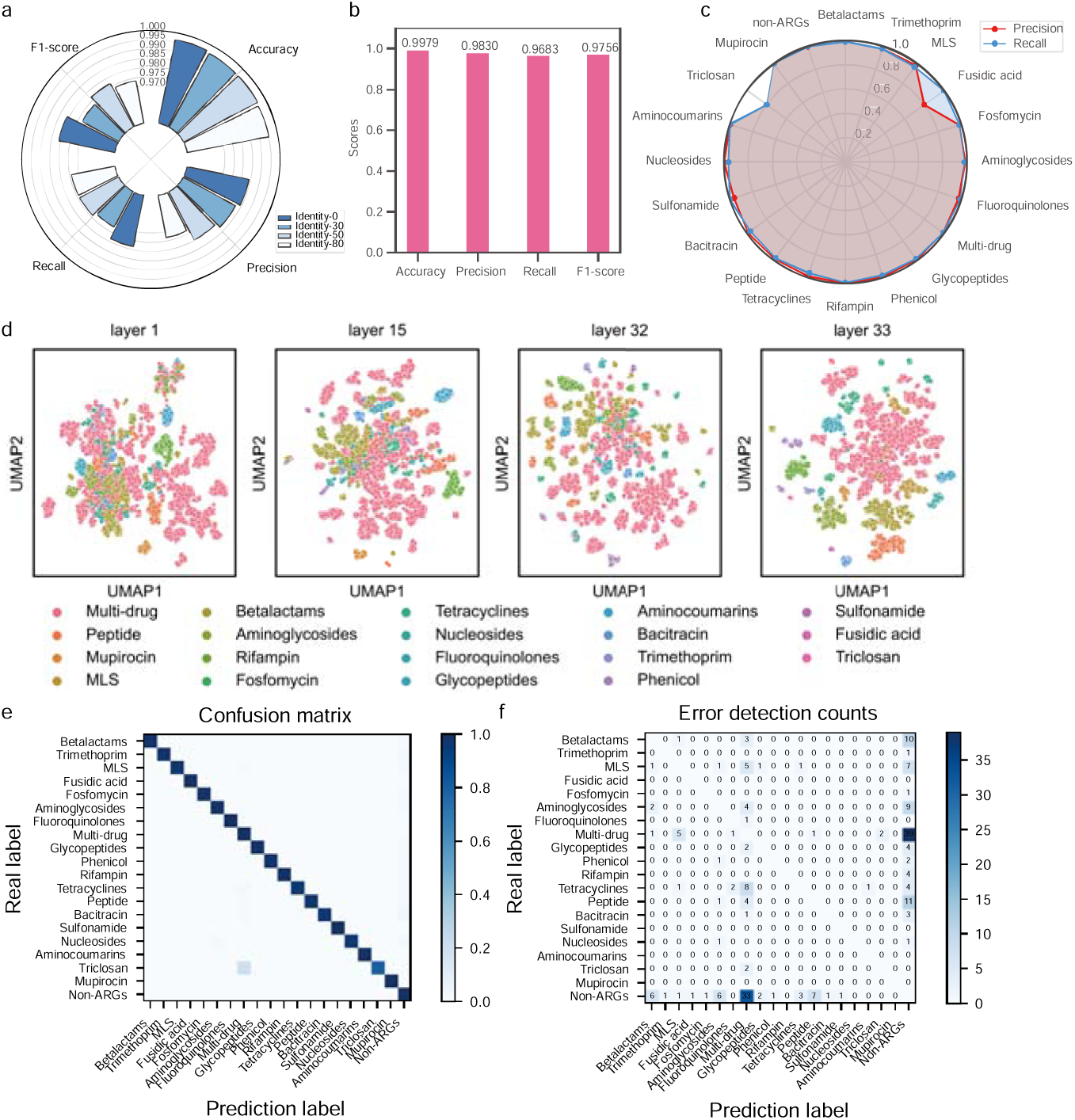
Performance evaluation of deep-learning FunGeneTyper models with structured Antibiotic Resistance Gene Database (SARD) for classification of ARGs. **a**, Evaluation of the influence of identity threshold used for selecting the negative dataset on model performance in the classification of ARGs. **b**, Performance metrics of ARGTyper developed based on FunGeneTyper models and SARD. **c**, Performance of all 19 classes as indicated by precision and recall of ARGs and non-ARG classes. **d**, Visualization of feature learning at different layers during the ARGTyper training process. **e**, Confusion matrix for ARG class classification, confusion between true (y-axis) and predicted (x-axis) ARGs. **f**, Number of ARG protein sequences annotated incorrectly. MLS: Macrolides, Lincosamides and Streptogramines.

To demonstrate the powerful utility of FunGeneTyper, we used the reference protein sequences in SARD to train two transformer models (FunTrans and FunRep) and developed them as a new deep-learning ARGTyper deep-learning classifier. We used the trained ARGTyper to classify the testing set to validate the performance of the ARGTyper. The overall ARGTyper performance metrics prove that FunGeneTyper provides an excellent and robust framework for gene classification. Specifically, the optimal FunTrans model at the ARG class level reached an accuracy of 0.9979, a precision of 0.9830, a recall rate of 0.9683, and an F1 score of 0.9756 (Fig. 2b). Moreover, the prediction precision and recall of all 17 ARG classes exceeded 0.96 (Fig. 2c), apart from fusidic acid and triclosan, which showed lower precision and recall because they have only 21 and 53 reference sequences, respectively, in SARD (Dataset S3). More training data helps the model learn more features. Nonetheless, the power of FunTrans to classify these temporarily less-represented classes of ARGs will improve as more functionally-verified sequences will be available for model training.

The vector space generated by FunGeneTyper was semantically rich and encoded structural, evolutionary, and functional information. To explain what our model intuitively learns, we obtained representations of all classes of ARG and non-ARG sequences in the training set. We used uniform manifold approximation and projection (UMAP) to reduce data dimensions in each layer to two. Visualizations performed in the four essential representative layers (1st, 5th, 32nd, and 33^rd^) revealed the learning process of the model (Fig. 2d). All ARG sequences were highly entangled at the first level of encoding input. However, they became increasingly separated as the transformer model got deeper. Each type of ARG undergoes a process from dispersion to aggregation. This finding verified that FunTrans can efficiently learn the representation features of sequences from raw input data with high entanglement.

Prediction multiclass confusion matrix was used to represent the effect of FunTrans on the learning features of each ARG class. The results indicated that the FunTrans model was excellent at predicting all ARG classes (Fig. 2e). We continued to locate significant classification errors in the ARG classes using error detection counts (Fig. 2f). Prediction error was concentrated in the multidrug class. Specifically, 33 non-ARG sequences were mispredicted as multidrug resistance, whereas 39 multidrug resistance protein sequences were mispredicted as non-ARG sequences. The poor prediction performance of these proteins is mainly due to their high structural differences and diverse biological functions that include roles other than multidrug resistance^30^, making it challenging for a DL model to effectively learn sufficient discriminative features in the absence of sufficient training data. Multidrug efflux pumps^30^ export antibiotics and other diverse extraneous substrates, including organic solvents, toxic heavy metals, and antimicrobials, and also fulfill other key biological functions such as biofilm formation, quorum sensing and survival and pathogenicity of bacteria^30^. Therefore, multidrug resistance proteins or efflux pumps were not seriously considered as ARGs^17, 31^ and we recommend excluding their sequences from ARG analysis unless they can be reliably or unambiguously assigned to resistance functions of certain antibiotic classes.

Once the FunTrans model was shown to be robust and accurate in identifying ARGs and classifying them into 19 classes, we trained FunRep, which conducted a more detailed lower-level classification of ARGs into 2972 groups (Dataset S3). FunRep achieved an overall prediction accuracy of 0.9023 for all ARG groups (Dataset S4). We used UMAP to visualize FunRep model’s learning process. UMAP was used to visualize the characteristics of the final layer of all classes except the Fusidic acid class (21 sequences, Dataset S3). UMAP showed that FunRep can cluster the features of each group in the main ARG classes, including beta-lactams (5909 sequences), Macrolides-Lincosamides-Streptogramines (MLS, 2317 sequences), aminoglycosides (3483 sequences), and glycopeptides (2037 sequences) (Supplementary Figure 3).

In summary, we demonstrated the application of the FunGeneTyper framework to develop ARGTyper as the first transformer-based ARG classifier trained from a customized structured ARG database (SARD). The performance metrics of the testing set show that FunTrans and FunRep can achieve highly accurate (accuracy=0.998) and robust (F1-score=0.976) identification of all known types (classes) and subtypes (groups) of ARGs in the authoritative CARD. Both the accuracy and robustness of FunGeneTyper models outperform previously published results from DeepARG (accuracy>0.97, F1-score>0.93 ^9^) and HMD-ARG (accuracy=0.935, F1-score=0.893 ^32^) on their own testing sets of ARGs.

### Model performance in the discovery of new genes

The ‘twilight zone’ of protein sequence alignment is a complex, long-standing problem plaguing protein function prediction^33, 34^, limiting the discovery of PCGs in the largely uncultured microbial world. In contrast to classic SA-based tools, DL-based models (FunRep and FunTrans) of the FunGeneTyper framework are designed with unique features and intrinsic advantages for predicting remote homologues of protein sequences with guaranteed accuracy and robustness, as previously demonstrated for ARG classification.

To compare FunGeneTyper’s ability to identify new PCGs with those of existing methodologies, we evaluated its ability of its DL-based models to discover remote homologues by predicting experimentally-confirmed protein sequences of new ARGs discovered from three representative habitats: human gut (n = 168)^35^, wastewater treatment plants (n = 77)^11^, and soil (n = 52)^36–39^. We computed the predictive performance of FunGeneTyper classifier for ARGs (ARGTyper) and compared it with that of three state-of-the-art tools: DL-based tools (HMD-ARG^32^ and DeepARG^9^), SA-based tools (RGI^7^), and HMM-based tools (Resfams^18^) (Table 1). FunGeneTyper had higher accuracy, precision, recall, and F1-score for predicting new ARGs compared with HMD-ARG^32^ and DeepARG^9^. The significant improvement was primarily attributed to our implementation of the protein semantic models (i.e., FunTrans and FunRep) in FunGeneTyper, which can learn more hidden features of protein sequences, especially the context information^19, 21^, compared with the traditional one-hot encoding algorithm and the convolutional neural network used by HMD-ARG^32^ and the multilayer perceptron used by DeepARG^9^. Moreover, the overall classification performance of FunGeneTyper, as benchmarked by the F1-score (0.5445 to 0.6948), was much higher than that of the classic SA-based methods (0.0556 to 0.6598) and HMM-based methods (0.2630 to 0.5224) (Table 1). Although RGI also achieved high accuracy (0.8830) in human intestinal data, its precision (0.4545), recall (0.3968), and F1-score (0.4195) were much lower than those of the FunTrans model (0.7500, 0.6642, and 0.6948, respectively) because many of the new ARG sequences tested here fell below the commonly applied stringent identity cutoffs (> 95% RGI). It is expected that when a strict one-size-fits-all filter cutoff is applied to the alignment results, many false-negatives would result, limiting the discovery of ARGs that show a more remote homology to database sequences. The superior performance of the FunGeneTyper classifier over existing tools in identifying new ARGs was further evident when comparative tests were performed using wastewater treatment plant (WWTP) or soil samples compared with human gut samples (Table 1). This indicates that FunGeneTyper has a greater capacity to predict functional genes in complex environmental samples. To further resolve the superior predictive performance of FunGeneTyper for remote homologues of functional genes over existing tools, we divided the ARGs data into lower homology (≤50% identity) and higher homology (≥50% identity) datasets (Supplementary Figure 4). FunGeneTyper not only consistently achieved better classification performance of higher homology ARGs in all three sample groups (WWTP, soil, and human gut), it also showed outstanding performance at accurately and sensitively predicting the function of remote homologous sequences (Dataset S5).

**Table 1.** Performance comparison between FunGeneTyper and other alternative bioinformatics tools for the discovery of experimentally med new ARGs. In total, 297 experimentally confirmed ARGs sequences of human gut^35^ (n = 168), WWTPs^11^ (n = 77), and soil^36–39^ (n = cteria were included in the comparative analysis which was performed under the default settings of each deep-learning (DL)-based, ce alignment (SA)-based or Hidden Markov Model (HMM)-based tool recommended by the developers.

| Tools | Human gut (n=168) |  |  |  | WWTP (n=77) |  |  |  | Soil (n=52) |  |  |  |
| --- | --- | --- | --- | --- | --- | --- | --- | --- | --- | --- | --- | --- |
|  | Accuracy | Precision | Recall | F1-score | Accuracy | Precision | Recall | F1-score | Accuracy | Precision | Recall | F1-score |
| *DL-based tools* |  |  |  |  |  |  |  |  |  |  |  |  |
| FunGeneTyper | 0.8512 | **0.7500** | **0.6642** | **0.6948** | **0.7273** | **0.7500** | **0.5403** | **0.6072** | **0.8269** | 0.5926 | **0.5529** | **0.5445** |
| HMD-ARG | 0.8452 | 0.6000 | 0.5230 | 0.5486 | 0.5714 | 0.7161 | 0.3877 | 0.4589 | 0.8077 | **0.6000** | 0.4560 | 0.5119 |
| DeepARG | 0.3512 | 0.6250 | 0.4720 | 0.5149 | 0.1688 | 0.5714 | 0.1682 | 0.2591 | 0.2885 | 0.3750 | 0.1057 | 0.1607 |
| *SA-based tools* |  |  |  |  |  |  |  |  |  |  |  |  |
| RGI | 0.3452 | 0.6250 | 0.4596 | 0.5065 | 0.0390 | 0.3750 | 0.0349 | 0.0632 | 0.1538 | 0.1250 | 0.0357 | 0.0556 |
| *HMM-based tools* |  |  |  |  |  |  |  |  |  |  |  |  |
| Resfams | **0.8830** | 0.4545 | 0.3968 | 0.4195 | 0.6234 | 0.6250 | 0.4736 | 0.5224 | 0.8088 | 0.2727 | 0.2545 | 0.2630 |

Taken together, our results exemplify the discovery and classification of novel ARGs, especially among relatively remote homologues (<50% identity), and demonstrate that FunGeneTyper is best at predicting new ARG protein sequences, exhibiting unprecedented capacity to identify new genes with high accuracy, sensitivity, and robustness.

### Evaluating the generalizability of FunGeneTyper

To demonstrate the generalizability of FunTrans and FunRep in classifying other gene categories, we trained the models using a calibrated and professionally expanded bacterial virulence factor database, VFNet^40^ and utilized them to develop a new transformer-based classifier of virulence factor gene (VFG), named VFGTyper. Before training the model, we built a two-level expert-curated structured database based on the virulence ontology and reference sequences in the VFNet database. Semantic and categorically ambiguous data were cleaned (Methods). The final structured virulence factor database (SVFD) consisted of 160484 VFG sequences distributed into 2837 classes in 45 families (Dataset S6).

We followed a strategy similar to that mentioned above to train the model, collecting a non-target dataset with 551,783 sequences (excluding VFGs) from Swiss-Prot, as the negative dataset (see Methods). With the merit of the proposed adapter module, we only need to re-train a new adapter when building a VFGTyper. The design of adapter allows us to train only a new classifier and an adapter when predicting new functions. All the parameters in the backbone network can be reused.

Therefore, VFGTyper can be regarded as a new task branch in the FunGeneTyper framework, where only the adapter and classifier differ. We verified the VFGTyper using the testing set to provide evidence of its generalizability in the functional genotyping process. VFGTyper achieved an accuracy of 0.9907 (Fig. 3a) in the family level prediction task. The obfuscation matrix results also showed that FunTrans achieved excellent classification performance for each VFG at the family level (Fig. 3b, Supplementary Figure 5). In addition, FunRep was 0.9499 accurate at predicting different VFG classes in the second-stage prediction.

**Fig. 3.**
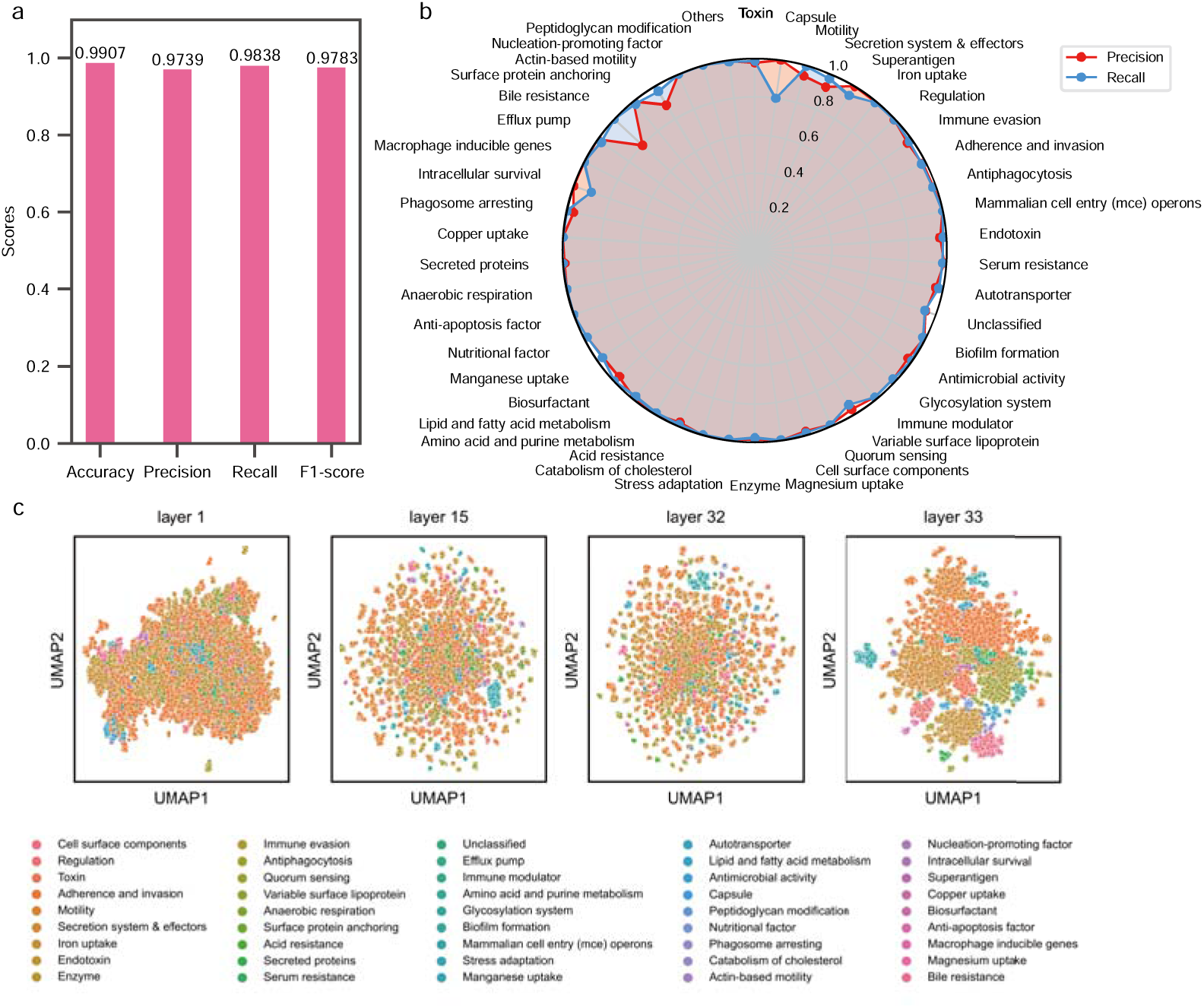
Transfer learning of FunGeneTyper models on Structured Virulence Factor Gene Database (VFGD) and performance evaluation for VFG classification. **a**, Performance metrics of VFGTyper developed based on FunGeneTyper models and VFGD. **b**, Precision and Recall of VFGs family and non-VFGs category. **c**, Visualization of feature learning at different layers in VFGs FunTrans training. VFGs: virulence factor genes.

In conclusion, we demonstrated that FunGeneTyper can be successfully generalized to develop VFGTyper as the first transformer-based VFG classifier of its kind and applied the new classifier to achieve ultra-accurate classification of VFGs by adding new adapters. To vividly show that FunGeneTyper can learn sufficient discriminative features from different groups of functional gene datasets, we visualized the learning process of FunTrans and FunRep models for VFG sequences. Consistent with the learning process for ARGs (Fig. 2d), both models also achieved effective feature clustering and classification of VFGs at both the family (Fig. 3c) and class (Supplementary Figure 6) levels. Besides classification performance, we also proved VFGTyper’s full capability in the discovery of an experimentally-confirmed novel VFG (NCBI accession no.: WP_034687872.1) of a toxin family in *Chryseobacterium piperi* with sequence similarity to botulinum neurotoxins (BoNTs) through re-analysis of published genome ^41^. Specifically, of the 8 putative toxin genes of *C. piperi* showing no significant (n=6) or only limited (n=2) sequence homology (i.e., global identity < 10%) to known reference VFGs, 7 were effectively identified as VFGs by FunGeneTyper and 4 were further classified as BoNTs (Dataset S7). Compared with a conventional sequence alignment (SA)-based approach which failed to predict 6 VFGs, the deep learning models of FunGeneTyper showed much greater capacity for the discovery of remote homologues of known toxin genes. Therefore, FunGeneTyper represents an expandable deep learning-based framework for ultra-accurate classification and discovery of functional genes, as demonstrated here for ARGs and VFGs.

### Privacy-preserving global sharing of plug-and-play adapters for functional gene discovery

To demonstrate the parameter efficiency of FunGeneTyper’s adapter modules, all 650 million parameters of the pre-trained model are fine-tuned as a benchmark test which achieved excellent prediction accuracy in ARGs class (0.9988) and VFGs family (0.9930). Comparatively, with only fine-tuning of about 21 million parameters (3% of all parameters) of the Adapter layer, we demonstrated that FunGeneTyper achieved near-identical excellent performance of 0.9979 for ARGs class and 0.9907 for VFGs family, proving that parameter-efficient lightweight plug-and-play adapter modules of FunGeneTyper can be easily shared without little loss of prediction accuracy.

Benefiting from the parameter-efficient property, FunGeneTyper has two novel merits. First, FunGeneTyper enables effective effort-sharing by the entire community (Fig. 4). Specifically, a researcher who has trained our FunGeneTyper model for classification or discovery of functional genes (other than ARGs and VFGs demonstrated here) can submit their adapters (along with a classification layer) to the adapter hub. Once the adapter has been submitted, the module can be downloaded and easily inserted into the FunGeneTyper model for direct downstream user application. Second, the adapter design helps solve data privacy issues. Where researchers have not publicly released their own datasets, they can train FunGeneTyper with their private datasets, submit only the adapter module (again along with a classification layer), and provide functional descriptions of their FunGeneTyper. Thus, the private datasets are protected, while the uploaded adapter models can be used without model training. As the number of researchers getting involved in the development of FunGeneTyper increases, the model may become a universal toolkit that can be used for predicting functional genes simply by looking up related functional modules. We believe that with the elegant adapter module, FunGeneTyper will facilitate adapter sharing and model integration globally.

**Fig. 4.**
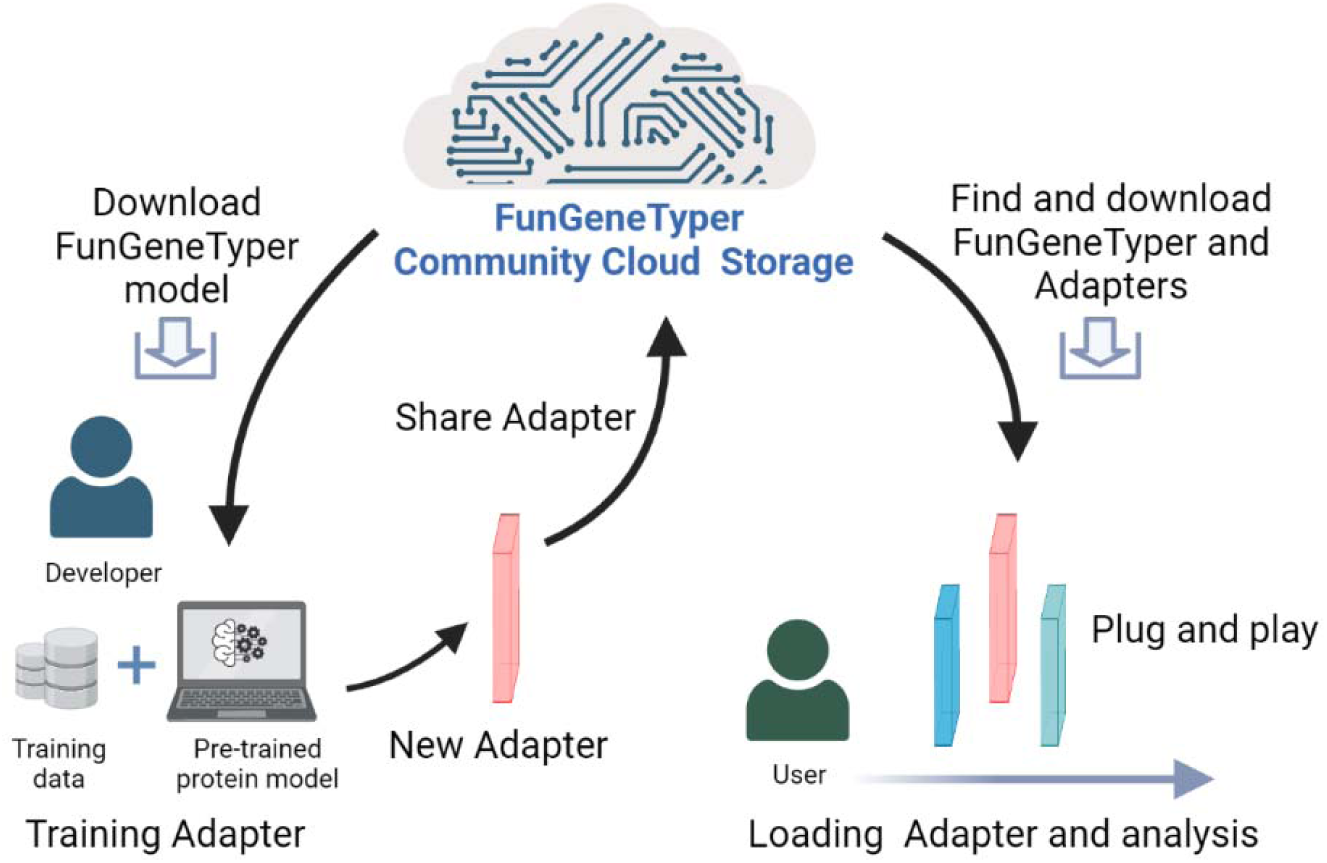
Schematic of the Adapter Sharing Community (ASC) in the framework of FunGeneTyper. The community developers are cyber de-centralized to train customizable structured databases and develop deep learning classifiers of various categories of functional genes, while users utilize the classifiers of interest to accelerate the discovery of genes which, in turn, provide new experimentally-confirmed sequences to expand the structured databases and improve deep-learning models.

## Discussion

Metagenomics has provided an opportunity for identifying microbiome diversity and novel functionalities. However, the speed at which high-throughput DNA sequencing technologies unravel the vast genetic novelties of uncultured microbes in nature outpaces our capacity to understand their function. Previous approaches for functional classification of PCGs were based on sequence alignment using tools such as BLAST^4^, usearch^5^, and Diamond^6^ or conserved motifs and domains using Hidden Markov Models. Selection of uniform cutoffs and thresholds usually limit the accuracy and/or sensitivity of these methods for functional gene prediction. Protein semantic algorithms based on NLP methods have been developed ^20, 24^. However, these algorithms are not optimized for classifying different categories of microbial genes, and a unified thinking paradigm is required to meet the needs for accelerated discovery of new genes.

Our study provides an expandable deep learning-based framework for efficient and robust gene function prediction, which represents an emerging methodological paradigm for global developers and users to tackle unprecedented challenges and meet the above-mentioned urgent needs in the classification and discovery of any group of functional PCGs. We propose an end-to-end FunGeneTyper framework for the classification prediction of gene functions. We exemplify the framework by developing two transformer-based classifiers, ARGTyper and VFGTyper, based on deep learning models coupled with expert-curated structured databases (SARD and SVFD) to realize robust functional classification of bacterial ARGs and VFGs, which are two categories of genes key to WHO’s one health approach for human, animal, and environmental health protection^42^.

A series of experimental validations, including five-fold cross-validation, testing set validation, and experimentally-confirmed protein sequence validation, demonstrate the effectiveness and robustness of FunGeneTyper. Using ARG as an example, FunGeneTyper models are more effective than SA-based and DL-based models in predicting protein sequences of new ARGs from the human gut, WWTP, and soil microbiomes with relatively low homology (< 50% similarity) to known ARGs. This shows that ARGTyper has an unmatched advantage in discovering ARGs, primarily because of the powerful learning ability of protein semantic models. Since experimentally-confirmed sequences of the major categories, types, and subtypes of genes are not sufficient, expanding the database based on sequence homology is common and necessary to obtain sufficient training sequence data. UMAP analysis showed that the expanded sequences represent reliable datasets and support our model to better learn discriminative protein semantic features to achieve satisfactory performance in identifying functional genes, including ARGs and VFGs.

Accurately classifying target genes from the huge interference of non-target gene data is a problem. Therefore, we purposefully introduced non-functional genetic datasets as part of the training set. Although this operation increases training complexity, it enables our model to accurately classify target genes from noisy data when used to analyze large-scale metagenomic sequence datasets from environmental or animal microbiome samples. Some machine learning methods rely on sequence alignment tools to create a similarity score matrix of potential gene sequences and databases^9, 43^. Such practices will inevitably be affected (and limited) by the selection of arbitrary thresholds for the results. The FunGeneTyper framework proposed here can accurately annotate genes via classification through discriminative features learned from multiple sequences. The limited number of training sequences may prevent the models from learning sufficient features. This transient issue would, however, be easily solved once more experimentally-confirmed reference protein sequences of target genes are available for model retraining and refinement. Meanwhile, the robustness of deep learning to noise labels^44^ can also help our framework models and classifiers outperform existing ones in discovering new genes. In particular, once large amounts of (meta)genomic data are freely available, a uniform and convenient understanding of the relationship between microbial gene sequences and protein function becomes a perennial challenge that can be tackled to create opportunities for gene discovery. There are other gene categories, apart from ARGs and VFGs, including those associated with microbially-driven global biogeochemical cycling (carbon, nitrogen, phosphorus, and sulfur) or microbial biodegradation (bioremediation and bio-restoration) and biosynthesis (biomedicine and bioresources) (Fig. 5), such as those well established by the RDP’s FunGene database^45^. Building a dynamic metagenomic bioinformatics community will help us better understand gene function. In principle, FunGeneTyper can predict the function of any gene category based on prior parameters of the pre-trained model and the adapter’s transfer-learning ability. The adapter module used in FunGeneTyper is a lightweight plug-and-play neural network that only fine-tunes and maintains a small set of parameters and is conducive for sharing and promotion. Therefore, other researchers can use the framework and training parameters we provide to train their own core datasets to easily develop predictive deep learning models of genes of interest. Researchers can also share a trained adapter through the adapter sharing community (ASC) without disclosing their private datasets. The future prosperity and collaboration of the ASC under the guidance of FunGeneTyper framework provide an interactive, dynamic, and continuously improving or evolving platform for functional classification of various PCG sequences. More importantly, FunGeneTyper and ASC are expected to contribute significantly to advances in industrial biotechnology, health and medicine, food and agriculture, environmental biotechnology, and bioenergy (Fig. 5), as they are increasingly applied to accelerate the discovery of new genes and enzymatic resources from microbiomes.

**Fig. 5.**
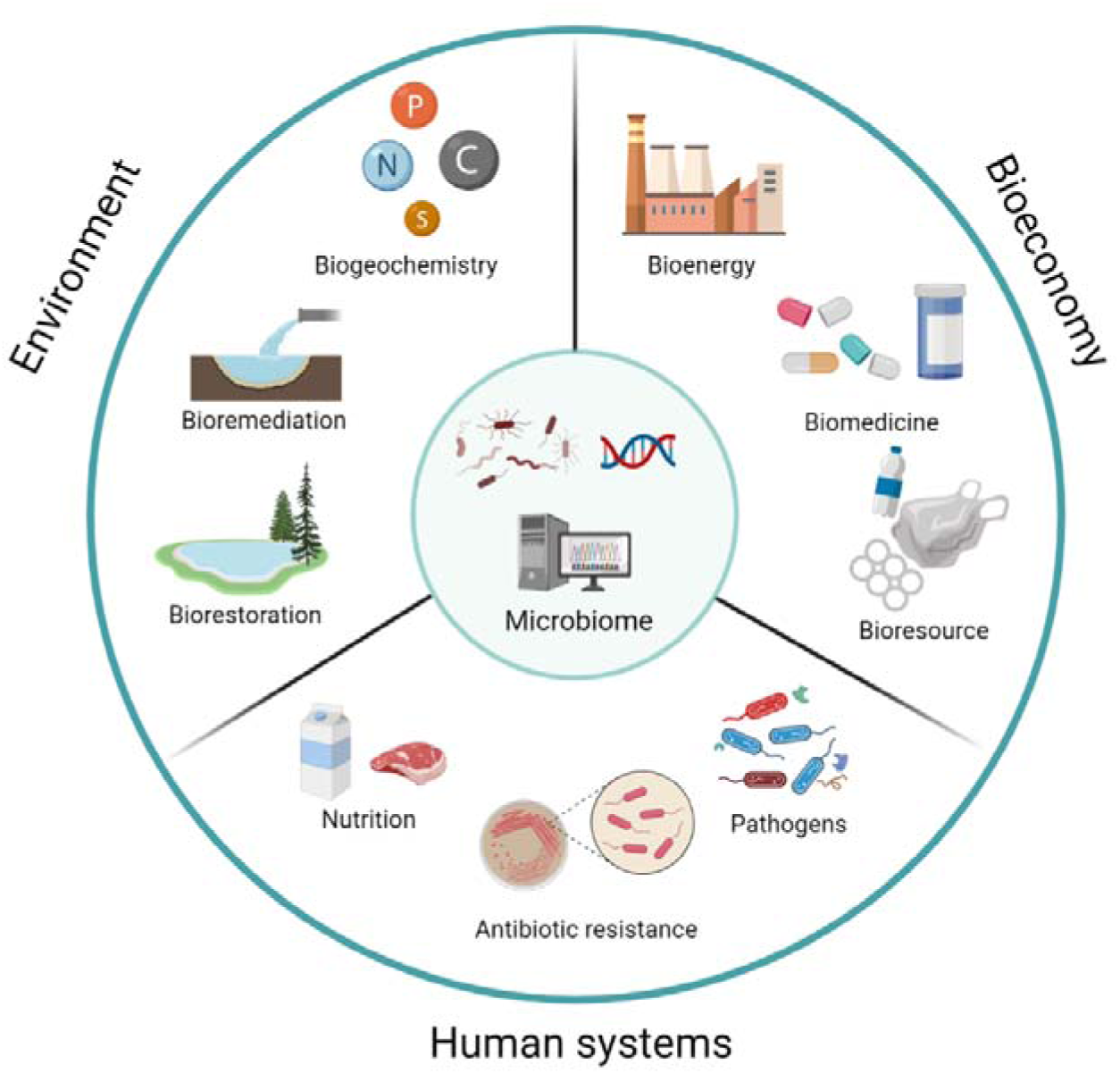
Potential applications of FunGeneTyper to the discovery of microbiome resources for enhancing our environment, bioeconomy, and human systems. Metagenomic discovery of precious genetic and enzymatic resources facilitated by the Adapter Sharing Community of FunGeneTyper can contribute to follow-up microbiome, genetic and protein engineering researches for enhancing human health and eco-environment systems.

In conclusion, FunGeneTyper provides an innovative and unified framework with deep learning models (i.e., FunTrans and FunRep), expandable classifier toolkits (e.g., ARGTyper and VFGTyper) and customizable structured databases for the ultra-accurate classification and discovery of functional genes (e.g., ARGs and VFGs) that have scientific and biotechnological significance. This framework will contribute to the robust monitoring of function genes and discovery of novel enzymatic resources from diverse microbiomes and uncultured microbes therein, which is critical to understand and harness the microbiome sciences underlying environment (biogeochemistry, bio-restoration, and bioremediation)^14^ bioeconomy (bioenergy and bioresources)^13^, and human systems (food and health)^20, 46^.

## Methods

### Collection and expansion of the core dataset

The core dataset used for FunGeneTyper model training is a set of experimentally-confirmed reference sequences of target functional genes collected from literature and/or expert-curated databases. Because the core dataset does not always contain a sufficient number of experimentally-confirmed sequences (no more than 10 sequences^40^) for every type or subtype of functional gene, it is expanded to retrieve more sequence data to improve and optimize the training of deep learning models. In the subsequent training method, which separates the extended categories of five or more sequences into the training set, verification set, and testing set at a ratio of 6:2:2, any categories that are unsuitable for inclusion in these five sequences are included in the training set.

### Construction of structured antibiotic resistance database (SARD)

#### Core ARGs dataset

To ensure the professionalism and accuracy of the training dataset, reference protein sequences of ARGs defined by homologs in the authoritative CARD were selected as core data for downstream model training. The sequences were clustered using CD-HIT^47^ (*v*4.8.1) at an amino acid sequence identity of 100%, and all protein sequences and their ontological information were manually checked to ensure that each ARG was properly classified into class (type) and group (subtype) based on their ontological information. Generally, class is equivalent to CARD’s ontology terms for antibiotic drug types, and group is equivalent to the specific sequence category. Macrolides, lincosamides, and streptogramins were combined into the MLS class. Based on the above procedures, a core dataset of 2972 non-redundant sequences representing 2972 groups of ARGs from 19 classes was obtained and used to build the SARD, which was used in subsequent analyses.

#### Expanded ARGs dataset

To ensure sufficient training data, the core dataset was expanded by retrieving close homologues of its ARGs from the Uniref100 database following strict screening criteria. Briefly, Diamond^6^ (version 2.0.15) was used to index the ARG sequences in the core dataset and to search for homologous sequences with an amino acid identity and coverage greater than or equal to 80%. The extracted candidate sequences were dereplicated and used as expanded datasets.

#### Negative dataset

To ensure that the model can learn sufficient features of non-target gene function, which is essential for robustly predicting target function directly from metagenomic data, we used the Swiss-Prot database, an expert-validated protein database, to generate a negative dataset for use as a non-ARG training set. First, protein sequences associated with antibiotic resistance in the Swiss-Prot database were screened out using the keywords KW-0046. The remaining sequences were aligned against the core ARGs dataset using Diamond software. Sequences with an alignment coverage greater than 80% were extracted and categorized into four negative datasets based on their sequence alignment identity (ID): identity 0 (ID ≤0%), identity 30 (ID ≤30%), identity 50 (ID ≤50%), and identity 80 (ID ≤80%).

### Construction of structured virulence factor database (SVFD)

#### Core VFGs dataset

Virulence factor databases were collected from VFNet^40^. Zheng et al^40^ performed a detailed similarity search for known and potential VFGs in the complete bacterial genome downloaded from the NCBI server using VFanalyzer^48^, with Virulence Factor Database (VFDB) as the core database^48^. VFNet is an expanded virulence factor database that can be used directly in the training process.

#### Negative dataset

The non-VFG collection process is similar to that of the non-ARG collection process, except that KW-0800 is used to filter sequences from Swiss-Prot database (version: June 2021).

### Architecture of the FunGeneTyper model

FunGeneTyper is a universal function classification framework composed of two core deep learning models, FunTrans and FunRep, which share similar structures but are designed to classify functional genes at the type and subtype levels, respectively. Both models are modular adapter-based architectures that leverage a few extra parameters to achieve efficient fine-tuning of large-scale PLMs. In detail, utilizing the state-of-the-art large-scale protein PLM esm-1b as a 33-layer transformer encoder framework as the foundation, we plug adapters in each transformer layer of the PLM, which are individual modular units that are used as newly introduced weights to be fine-tuned for specific functional tasks. Notably, ESM-1b, through self-supervised learning on the UniRef50 dataset, was shown to have a superior capacity to infer fundamental structural and functional characteristics of proteins from gene sequences^49^.

The holistic architecture is depicted in Fig. 1a and consists of three main components: a multi-headed self-attention, a feed-forward network, and an adapter layer. Each sublayer contains layer normalization and skip connections to effectively train the neural network and avoid overfitting. It is worth noting that the bottleneck-shaped adapter module consists of a down-project linear *H E LJ^dXk^H E ^dXk^* where *d* is embedding size of the Transformer model, *k* is the dimension of the adapter and d» *k*, a ReLU activation followed by an up-projection *LE LJ^kXd^ LE LJ^kXd^*. The adapter layer is formulated as follows:

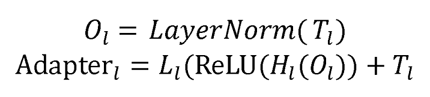

where *T_l_* is the hidden feature at transformer layer *l, d* = 1280, and *k* = 256 in the actual training.

Following the approach of BERT^50^, hidden features from the first token of the sequence of the last layer are extracted. In contrast to FunTrans, which adds a nonlinear layer for protein function classification after the representations of the last layer, FunRep first computes the hidden features of experimentally-confirmed core sequences and then annotates PCGs by finding the sequence’s category with the closest Euclidean distance in the representation space.

Here, we use a dual-tower architecture with shared parameters similar to Sentence-BERT^51^ for model training in order to place sequences with the same category closer in the representation space. FunRep is trained by constructing *< A,P,N >* triples, where *A* is the anchor sequence, *P* is a positive example possessing the same category as *A*, and *N*is a negative example whose category is different from A and the hidden representations they obtained through FunRep area, *p*, and *n*, respectively. The loss function adopts Triplet Loss, which is defined as follows:

*Loss(a,p,n) = max(D(a,p)-D(a,n)) + margin 0)*

where *D* is the Euclidean distance between vectors, and margin is an adjustable threshold, set to 1.0 during model training. ARGTyper-FunRep and VFGTyper-FunRep are classified at the group level with the same 21.76M learnable training parameters.

### Training settings

All datasets are divided into training, validation and testing in a 6:2:2 ratio, and the five-fold cross-validation is performed. Adam optimizer with default parameters is used, dropout is set to 0.2, learning rate is 1e-5, and the early stopping method is adopted to prevent overfitting. The accuracy, precision, recall and F1-score are used to evaluate the performance. As a result, the micro average of the F1-score also equals that of precision and recall, as well as the overall accuracy. Thus, we report only the overall accuracy for the micro average metrics while reporting precision, recall and F1-score for the macro average metrics.

### Evaluation of FunGeneTyper for the discovery of new functional genes

To validate the capacity of FunGeneTyper models in discovering new functional genes, experimentally confirmed ARGs from functional metagenomics studies were retrieved from NCBI’s protein database (accession numbers in Dataset S8). After removing those ARGs showing perfect sequence match to the CARD database (or core dataset of ARGs), 297 experimentally confirmed ARG sequences of human gut^35^ (n = 168), WWTPs^11^ (n = 77), and soil^36–39^ (n = 52) bacteria were retained for use in the downstream comparisons between FunGeneTyper and the well-established SA-based (RGI^7^), HMM-based (Resfams^18^), and DL-based (DeepARG^9^ and HMD-ARG^32^) approaches in terms of classification performance of the new ARGs. In this study, four evaluation metrics including the accuracy, precision, recall and F1-score were computed to assess the multi-classification results performance using the following equations:

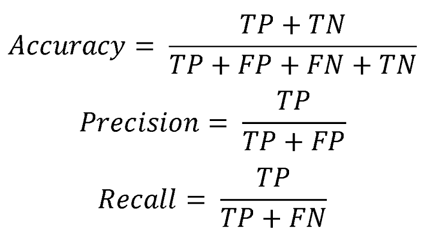

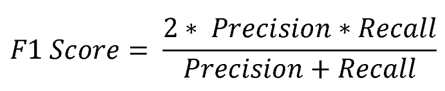

where TP is the number of true positives, TN is the number of true negatives, FP is the number of false positives, and FN is the number of false negatives.

To compare the ability of FunGeneTyper for discovering new VFGs, BoNTs-like sequences from the genome of *Chryseobacterium piperi* reported in a prior study ^41^ was downloaded from NCBI’s database by accession number (Dataset S7). Then, VFGTyper was used to predict VFGs and their affiliated family from the BoNTs-like sequences, and the output results were compared with those by a conventional sequence alignment-based approach with Diamond^6^ (version 2.0.15) search of the BoNTs-like sequences against SVFD.

## Abbreviations

DL: Deep Learning
SFGD: Structured Functional Gene Datase
PCG: Protein-Coding Gene
ARG: Antibiotic Resistance Gene
VFG: Virulence Factor Gene
MLS: Macrolides, Lincosamides and Streptogramines

## Author Contributions

1. F. Ju conceived the FunGeneTyper framework idea, obtained funding, and supervised the project. F. Yuan designed the Adapter sharing mechanism. G. Zhang and H. Wang (visiting student from Northeastern University) performed the model construction, data analysis and visualization. F. Ju and F. Yuan co-supervised G. Zhang and H. Wang on the deep-learning model construction with additional support from G. Guo. G. Zhang built the structured databases and accomplished data presentation with the assistance from J. Yang, Z. Zhang, L. Zhang. F. Ju and G. Zhang co-wrote the manuscript with assistance from F. Yuan and H. Wang. J. All authors approved the final version of the manuscript.

## Acknowledgement

The authors would like to thank Xinyu Huang and Lingrong Jin for valuable discussion.

We thank Yisong Xu for her professional support in lab management. We thank the Westlake University High-Performance Computing Center for computation support. We thank Kangyong Hu, Ling Yang, and Hang Li for their support of server maintenance.

## Footnotes

* The codes and database resources are available at: https://github.com/emblab-westlake/FunGeneTyper.

