## Supplementary Information for "Ultra-Accurate Classification and Discovery of Functional Protein-Coding Genes from Microbiomes Using FunGeneTyper: An Expandable Deep Learning-Based Framework"

**Figure and Table Legends**

**Supplementary Figure 1** The identity threshold for negative dataset expansion.

**Supplementary Figure 2** The number of ARGs class sequences in the core SARD.

**Supplementary Figure 3** Visualization of feature learning at the layer 33 in ARGs FunRep training. Each point is a two-dimensional mapping of a sequence of higher-dimensional representations.

**Supplementary Figure 4** An overview of the alignment of experimentally confirmed protein sequences.

**Supplementary Figure 5** Confusion matrix for VFs family level annotations.

**Supplementary Figure 6** Visualization of feature learning at the layer 33 in FunRep training of VFs.

**Dataset S1.** The details of ARGs in the core dataset of Structured Antibiotic Resistance Database (SARD). The core dataset of SARD is originated from the reference protein sequences of ARGs defined in the authoritative comprehensive antibiotic resistance database (CARD). The naming of Gene family, Subclass and Group is according to the Antibiotic Resistance Ontology (ARO) information in the CARD was manually inspected.

**Dataset S2.** The details of ARGs in the core and expanded datasets of Structured Antibiotic Resistance Database (SARD). The SARD includes 2972 core ARGs from CARD and 58902 expanded ARGs from Uniref100.

**Dataset S3.** The count statistics of ARGs class and group in the Structured Antibiotic Resistance Database (SARD). The ARGs were classified into 20 classes and 2972 groups.

**Dataset S4.** The classification performance of ARGTyper on the test set of ARGs from Structured Antibiotic Resistance Database (SARD).

**Dataset S5.** Comparison of classification results in experimentally-confirmed protein sequences. FunGeneTyper models showed outstanding performance in predicting the function of remote homologous sequences (identity < 50%) with greater accuracy and sensitivity than existing tools.

**Dataset S6.** The count statistics of family and class of virulence factor genes (VFGs) in the structure virulence factor dataset (SVFD). The VFGs in SVFD were classified into 45 families and 2837 classes.

**Dataset S7.** FunGeneTyper showed greater capacity for the discovery of remote homologues of known toxin genes compared with sequence alignment (SA)-based approach. Among the 8 putative toxin genes of C. *piperi* showing no significant (n=6) or only limited (n=2) sequence homology (i.e., global identity < 10%) to matched reference VFGs in SVFD, 7 were effectively identified as VFGs by FunGeneTyper and 4 were further classified as BoNTs. However, the conventional sequence alignment (SA)-based approach based on Diamond research failed to predict 6 of the 8 putative VFGs.

**Dataset S8.** The list of experimentally confirmed protein sequences from Human Gut, WWTP and soil samples. This dataset was downloaded from the NCBI's database according to the Accession number, and those new ARG sequences not included in the reference database were used for comparative performance evaluation of FunGeneTyper with other bioinformatics tools in the discovery of new ARGs.

### Supplementary Figures

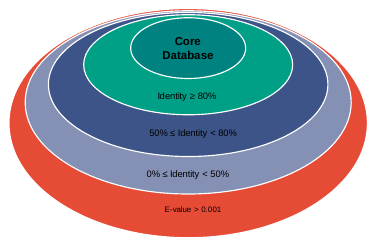

**Supplementary Figure 1** The identity threshold for negative dataset expansion. Four non-target sequence sets collected from four progressively stricter identity thresholds as the negative datasets.

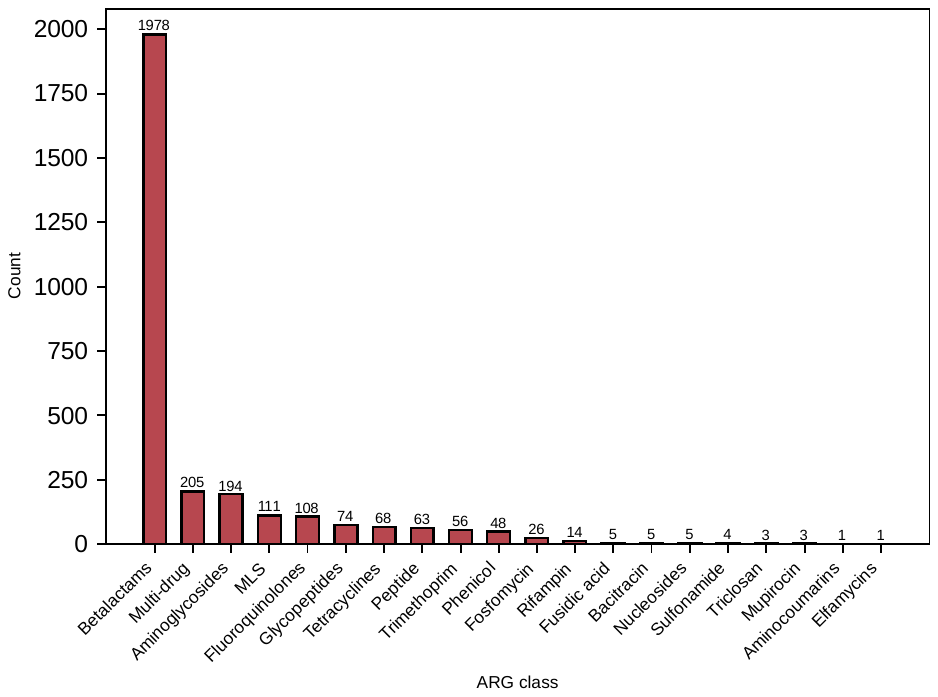

**Supplementary Figure 2** The number of ARGs class sequences in the core SARD. The x-axis is ARG class in core SARD. The y-axis is number of ARGs class sequences in the core SARD. SARD: structured ARG database.

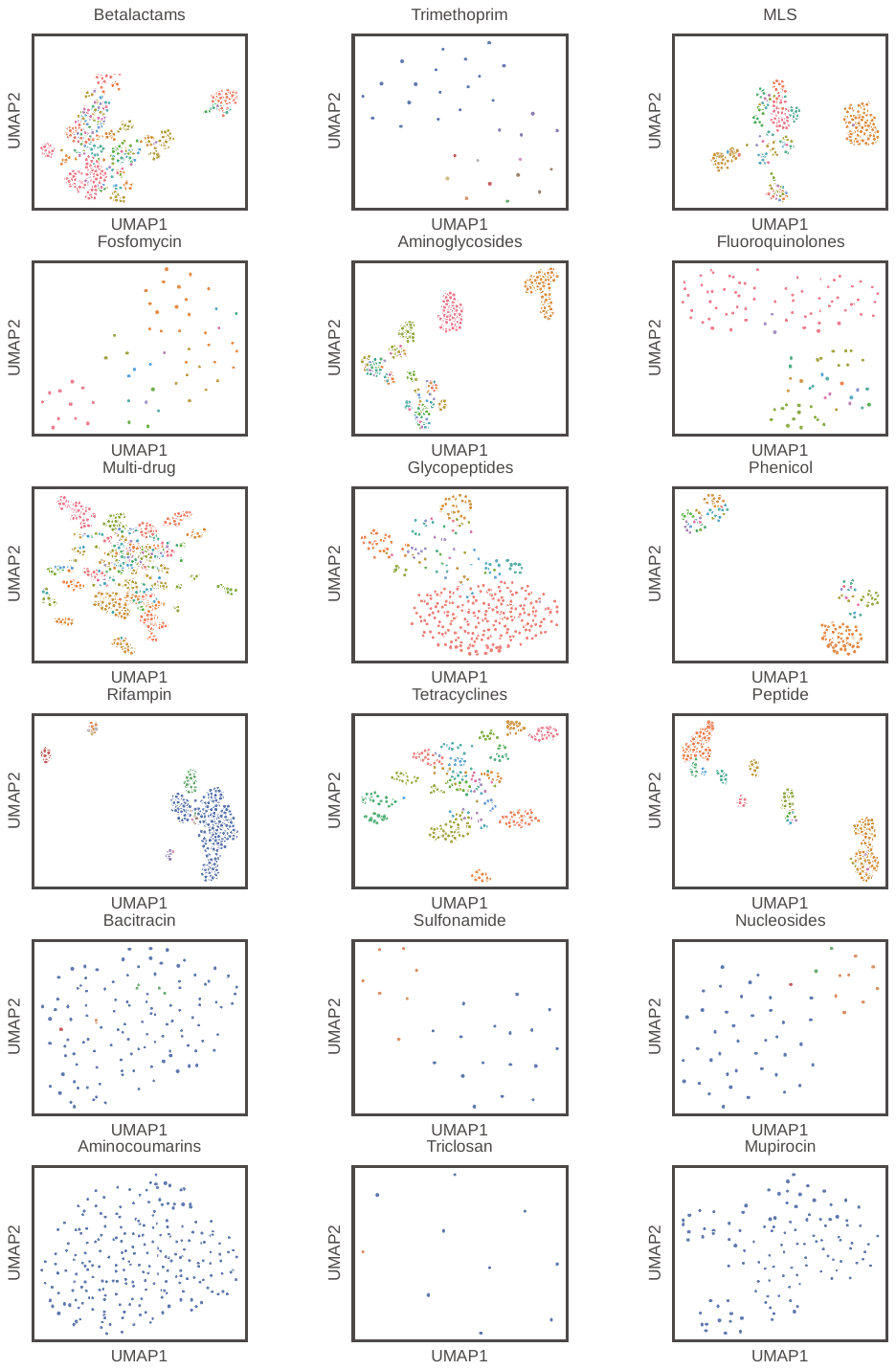

**Supplementary Figure 3** Visualization of feature learning at the layer 33 in ARGs FunRep training. Each point is a two-dimensional mapping of a sequence of higher-dimensional representations. The same color indicatesthe same ARGs group. UMAP analysis shows the clustering of features in the same ARGs group.

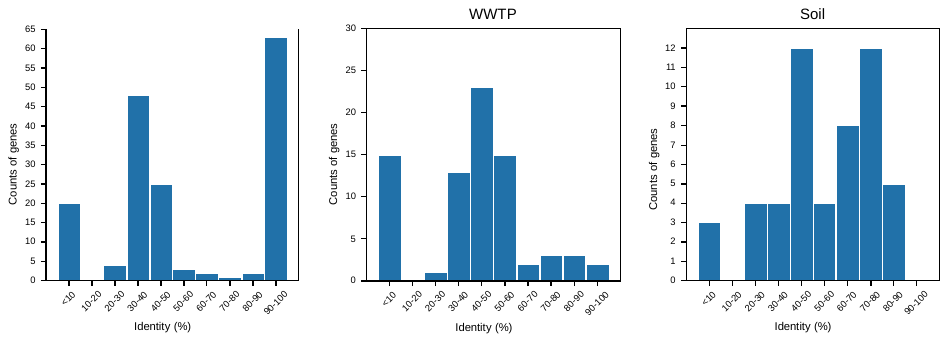

**Supplementary Figure 4** An overview of the alignment of experimentally confirmed protein sequences. The identity distribution of alignment between the experimentally confirmed sequence and the core ARGs dataset. Experimentally confirmed protein sequences was divided into relatively lower homology (≤50% identity) and relatively higher homology (≥50% identity) datasets.

**Supplementary Figure 5** Confusion matrix for VFGs family level annotations. Confusion matrix for VFGs family classification, confusion between true (y-axis) and predicted (x-axis) VFGs. VFGs: virulence factor genes.

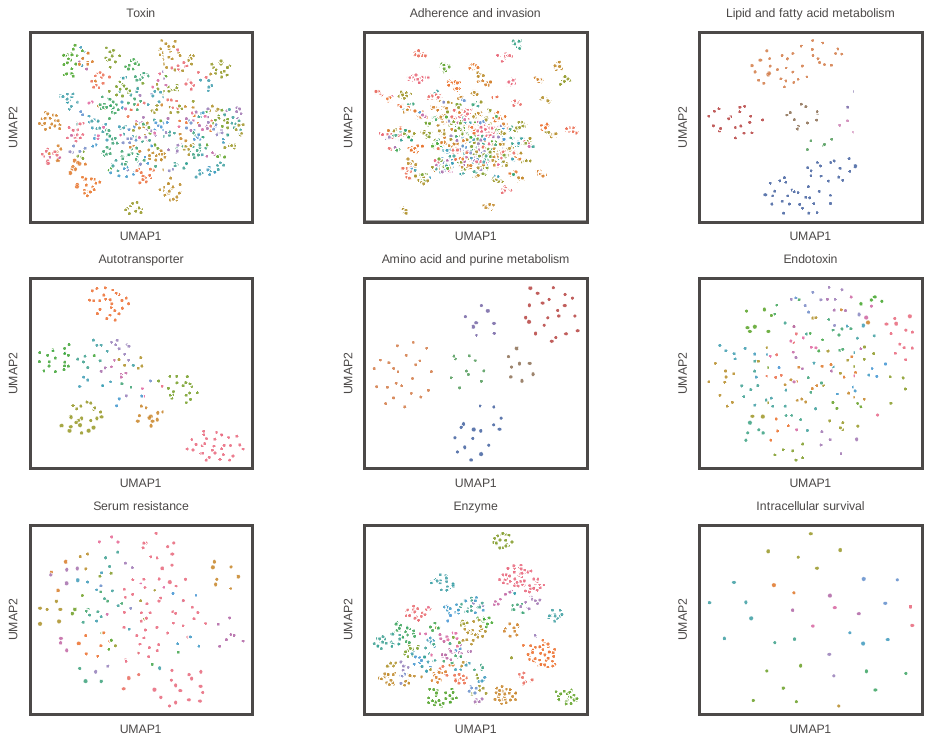

**Supplementary Figure** 6 Visualization of feature learning at the layer 33 in FunRep training of VFGs. Each point is a two-dimensional mapping of a sequence of higher-dimensional representations. The same color indicates the same VFGs class. UMAP analysis shows the clustering of features in the same VFGs class.
